# Benchmark of lasso-like penalties in the Cox model for TCGA datasets reveal improved performance with pre-filtering and wide differences between cancers

**DOI:** 10.1101/2020.03.09.984070

**Authors:** Rémy Jardillier, Florent Chatelain, Laurent Guyon

## Abstract

**Motivation:** Prediction of patient survival from tumor molecular ‘omics’ data is a key step toward personalized medicine. With this aim, the databases available are growing, with the collection of various ‘omics’ characterizations of patient tumors, together with their associated clinical outcomes for weeks to years of follow-up. Cox models with variable selection used with RNA profiling datasets are popular for identification of prognostic biomarkers and for clinical predictions. However, these models are confronted with the ‘curse of dimensionality’, as the number *p* of covariates (genes) can greatly exceed the number *n* of patients. To tackle this problem, variance-based pre-filtering and penalization methods are popular for dimension reduction. In the present paper, we study the impact of a pre-filtering step based on gene variability, and we evaluate the performance of the lasso penalization of the Cox model and four variants (i.e., elastic net, adaptive elastic net, ridge, univariate Cox) in terms of prediction, selection and stability.

**Results:** First, we show that the prediction capacity with the Cox penalties method is cancer dependent. Second, we develop a methodology to fix a threshold to filter out genes with low variability without losing prediction capacity. Third, we show that it is best not to use the Cox model to select prognostic biomarkers, as its false discovery proportion is always ≥ 50%. Finally, to predict overall survival, we can suggest the use of the ridge penalty, or the elastic net if a more parsimonious model is needed, after the pre-filtering step.

**Availability:** We provide the R script generated to reproduce all of the figures presented in this article.

**Supplementary information:** Supplementary Figures and R scripts are available.

## 1 Introduction

The roots of the ‘P4’ model of cancer medication lie in prediction combined with personalization, prevention, and participation [15]. Prediction of the best treatment for a given patient and prediction of various clinical outcomes, including overall survival, are both of growing interest. ‘Omics’ technologies now come with decreasing costs, which has made possible the molecular characterization of tumor samples of various types, including quantification of mutations and gene expression [29, 32]. As a result, there are growing numbers of knowledge databases that include molecular profiling of patient tumors, together with clinical information from patient follow-up. Survival analysis from transcriptome profiling of cancer patients in terms of messenger RNA (mRNA) expression is now emerging for clinical use [9].

The Cox proportional hazard model [6] is one of the most popular approaches in medicine to link covariates to survival data. [38] and [18] showed that Cox model are at least as good as, or even better than, neural networks and other machine learning models. Cox regression models with lasso penalty for variable selection [35] are often used to identify a few prognostic biomarkers from among the thousands of genes profiled, and to obtain a parsimonious model for simpler and cheaper clinical applications. When considering the numbers of covariates, *p* (which can typically be 20,000 gene products), in relation to the number of patients in the databases, *n* (which can typically be only a few hundred), various issues occur due to the high dimensionality, which include the lack of stability of the selected genes [16] and over-fitting [25]. This *p* ≫ *n* problem is referred to as “the curse of dimensionality” [1].

Lasso generalizations have been proposed for generalized linear models, such as Cox regression, to improve the performance and stability. In particular, the elastic net [40] and the adaptive elastic net [41] are regularization procedures that can overcome some stability issues of the lasso in the presence of highly correlated variables [39]. This occurs in particular for gene signatures. The adaptive elastic net also ensures additional theoretical properties to recover true biomarkers. As lasso-like methods, the ridge penalty allows control of the variance of the estimator. Although there is no selection, the ridge regression has been shown to have promise for reliable survival predictions with high-dimensional microarray data [36]. [24] showed that the lasso, ridge and elastic net penalties perform equally for low dimensional settings with few events. [3] compared the lasso, elastic net, ridge and adaptive lasso penalties on simulated and real acute myeloid leukemia data, and on real breast cancer data. They used the integrated Brier score as the metric to judge the performance of the models, and the datasets they used comprised fewer than 300 patients.

Pre-filtering procedures are popular methods that considerably reduce dataset dimensionality to avoid the curse of dimensionality issues. For instance, the less variable genes are subject to measurement noise and provide poor contributions to distinguish between patients [23]. Moreover, pre-filtering the less variable genes has been shown to increase the identification power of differentially expressed genes [13, 4]. Finally, this pre-filtering step is often done without justification of the threshold [11, 21, 22], while this value has been shown to be crucial to optimize the selection process [4].

Univariate selection procedures are also widely used methodologies to select features. [28] showed that univariate Cox regression is the best from among a set of eight univariate methods to select genes related to survival.

Finally, determination of the cohort size is a key issue to design any clinical trial [34]. If too few patients are included, inference procedures can provide non-significant results. On the other hand, including many patients is expensive, time-consuming, and requires more effort. Therefore, we investigated the impact of the number of patients on the predictive accuracy of the models.

To the best of our knowledge, no independent benchmark of univariate and multivariate (e.g., lasso, elastic net, ridge, adaptive elastic net) Cox regression has been defined using The Cancer Genome Atlas (TCGA) datasets, with pre-filtering of the genes based on their variability, and according to well-established metrics. In the context of survival analysis using high-dimension mRNA-seq datasets, the goals of this paper are to: (i) compare four multivariate Cox penalty methods (i.e., lasso, elastic net, adaptive elastic net, ridge) with univariate Cox regression and to each other; (ii) study the impact of pre-filtering the genes based on their interquartile range (IQR), and propose a rationale toward the fixing of a threshold; and (iii) study the impact of cohort size on the prediction performances. The methodologies are assessed in terms of the quality of the prediction of overall survival and prognostic biomarker selection, along with the stability.

To evaluate the Cox regression methods for prediction of overall survival and selection of genes correlated to survival, we use a panel of four cancers from TCGA (http://cancergenome.nih.gov/) that share different characteristics. Real datasets are used to study the prediction performances, and we provide realistic simulated datasets that maintain the complex structure of the molecular profiles of the patients to assess selection accuracy.

## 2 Materials and Methods

### 2.1 Cox proportional hazards model: the link between genetic and survival data

Let *T* denote the survival time (also called the ‘time-to-event’). The Cox model [6] is widely used in medicine to link covariates to survival data through the hazard function 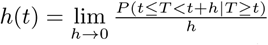, for all time instant *t >* 0, which represents the instantaneous death probability per unit of time. In the Cox model, the hazard function is defined as follows:

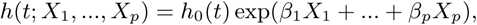

where *h*_0_(*t*) is the baseline hazard function, *X*_*j*_ are the covariates (here as gene expression), and *β*_*j*_ are the associated coefficients, for *j* = 1, *…, p*. For each patient *i* = 1, *…, n*, let *t*_*i*_ be the follow-up time (as either patient survival or censoring time), and *δ*_*i*_ the associated status, as 1 for death and 0 for censoring. The vector ((*t*_1_, *δ*_1_), *…*, (*t*_*n*_, *δ*_*n*_)) is referred to as the ‘survival data’. Let 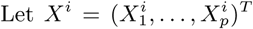 be the vector of covariates for the *i*−th patient. The vector of coefficients *β* = (*β*_1_, *…, β*_*p*_)^T^ can be estimated by maximizing the Cox pseudo-likelihood, as proposed by Breslow [5]:

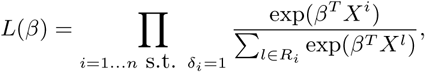

where for each patient *i* = 1, *…, n* whose death is observed (i.e., *δ*_*i*_ = 1), *R*_*i*_ is the set of patients at risk at time *t*_*i*_; i.e., such that *t*_*j*_ ≥ *t*_*i*_ (including patient *i*).

This function is called the ‘pseudo-likelihood’, because it is not a product of density functions, but a product of conditional probabilities. 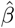 is computed by maximizing this pseudo-likelihood function: 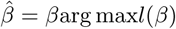, with *l*(β) = log(*L(*β)), the log-pseudo-likelihood.

Note that the Cox model is not intuitive, in the sense that it links genetic data to patient survival in an indirect way, through the hazard function. However, Cox pseudo-likelihood allows censored data to be efficiently dealt with. Moreover, this yields a robust inference procedure where the baseline function *h*_0_(*t*) does not need to be modeled or estimated in a parametric way. Finally, the estimation procedure leads to a convex optimization problem, for which efficient procedures and packages exist for computing 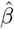 [12].

### 2.2 Penalization methods and univariate Cox selection for a sparse model

The main goal of survival analysis is to predict survival times with a set of covariates *X*. All of the 20,000 genes included do not necessarily contribute to the prediction of survival, and selection procedures are useful to reduce the dimension while retaining the most informative covariates.

For variable selection, we considered three alternative penalties in the likelihood of the Cox model: the lasso [35], the elastic net [40], and the adaptive elastic net [41] (implementation is provided in R script *functions/learn models func.R*). The adaptive elastic net uses a two-step procedure that includes the ridge regression [36]. Briefly, these methods consist of the addition of a penalty term to the log-pseudo-likelihood *l* before the maximization:

- The lasso

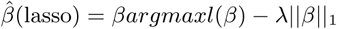
- The elastic net

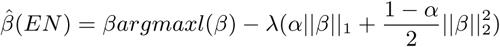
- The ridge

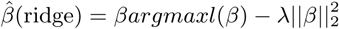
- The adaptive elastic net (a two-step procedure)
  1. estimates the vector 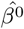 by maximizing *l* with the ridge regression.
  2. weights the elastic net penalty with the coefficient 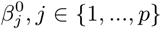 computed in step 1:

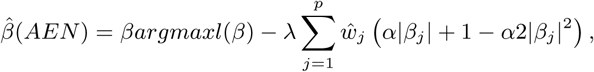

with 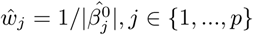, *j* ∈ {1, *…, p*}.

The *ℓ*_1_ norm forces some coefficient estimates 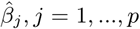, to be zero, and allows the selection to be made. For multivariate Cox selection models, genes selected are defined as the genes with nonzero 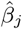 coefficients. It has been empirically observed that if there are high correlations between predictors, the ridge penalty provides better prediction performances than the lasso [35], and that the shrinkage effect of the lasso is too strong for large effects [39]. The elastic net and adaptive elastic net penalties have been developed to tackle these two issues, respectively.

The weight of the penalty, *λ*, is computed by K-fold cross-validation (K=5) using the package *glmnet* [12] in R version 3.6.0 [27]. The weight *λ* that minimizes deviation in the cross-validation is given by *λ*_*min*_. For more details on the mathematical concepts used in this article, we refer the reader to the book ‘The Statistical Analysis of Failure Time Data’ [17].

Univariate Cox selection consists of the learning of a Cox model individually for each gene. A p-value that corresponds to the statistical test with null hypothesis “*β*_*j*_ = 0” is then assigned to each gene *j* with 1 ≤ *j* ≤ *p*.

### 2.3 Filtering by interquartile-range

The Interquartile-Range (IQR) is a robust measurement of statistical dispersion, and it is defined as the difference between the 75th and 25th percentiles. We removed genes for which the IQR of expression among all the patients was below a given threshold, thus reducing the dimension. Bourgon et al. (2010) and Hackstad et al. (2009) showed that pre-filtering by variance increases the selection power of differentially expressed genes in high dimension analyses, and both of these studies indicated the crucial issue of the choice of the threshold. Additionally, the most variable genes present a better signal-to-noise ratio with respect to measurement noise. These genes are then easier to measure and more reliable, technically speaking.

### 2.4 Assessing the concordance index for prediction quality

The concordance index (C-index) allows the predictive ability of a risk score to be assessed by quantifying the proportion of comparable patient pairs whose risk scores are in good agreement with their survival data. For two patients *i* and *j* with risk scores *R*_*i*_ and *R*_*j*_, and with survival times *T*_*i*_ and *T*_*j*_, the C-index is defined as *C* = *P* (*T*_*i*_ *< T*_*j*_ | *R*_*i*_ *> R*_*j*_).

A C-index of 1 indicates perfect agreement, which means that the survival time can be perfectly described by a deterministic monotonically decreasing transformation of the risk score. Conversely, a C-index of 12 corresponds to random chance agreement, which means that the risk score do not provide any information on survival time. The risk prediction score used for multivariate Cox regressions is the Prognostic Index (PI) of the Cox model, which is classically defined as 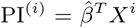 for patient *i*, 1 ≤ *i* ≤ *n*. These PIs are linear combinations of gene signatures for each patient that are computed in a testing dataset (as a random selection of 20% of the patients), with 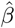 coefficients estimated with the Cox model in a training dataset (as 80% of the remaining patients). This procedure is repeated 100 times, and boxplots of the C-indices can be drawn for all of the methodologies and for different IQR thresholds. As the 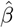 estimated are different for each penalization method, the PIs and the C-index are different for each method. For univariate Cox selection, the gene with the lowest p-value is selected from the training dataset, and we use the expression level of this gene as the risk prediction score to compute the C-index for the testing dataset (R script *prediction_quality.R*).

We took the estimator of the C-index given by [14] and theorized by [26], which is implemented in the package *survcomp* [31]:

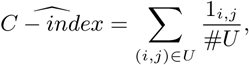

where *U* is the union of all of the comparable pairs (the death of the patient with the lowest follow-up time has to be observed), #*U* is the size of U, and 1_*i,j*_ is equal 1 if risk scores and survival data of patients i and j are in good agreement, and 0 otherwise.

As a comparison, we calculate the C-index estimates with the ridge regression under the null hypothesis “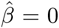”. This is done with the following permutation test procedure: the survival times are randomly shuffled, while the gene expression profiles are not modified, and the C-index is computed as explained in the previous paragraph. This procedure is repeated 100 times (i.e., 100 Monte-Carlo runs) to obtain the empirical null distribution of the C-index. This allows determination of whether the prediction performance, as computed in the previous paragraph, is significantly better than a purely random model.

### 2.5 The Cancer Genome Atlas dataset

First, in a preliminary study, we included all of the 11 cancers for which more than 500 patients with mRNA-seq data were available in TCGA (http://cancergenome.nih.gov/) (R script *cancer characteristics.R*). We studied the ‘time-to-death’, also called ‘overall survival’, which was defined as the time between diagnosis and death. We computed multiple characteristics for these 11 cancers, including the C-index estimates computed with the ridge regression method applied over all of the genes (Fig. 1). Then, we selected four cancers with different characteristics (Table 1) as a representative panel to evaluate the five methods (i.e., lasso, elastic net, adaptive elastic net, ridge, univariate Cox). We chose brain lower grade glioma (LGG) for its high C-index (0.81), breast invasive carcinoma (BRCA) because many samples were available (1040), and kidney renal clear cell carcinoma (KIRC) and lung squamous cell carcinoma (LUSC) as they had similar numbers of patients (524, 474, respectively) and censoring rates (0.69, 0.67, respectively), but very different C-indices (0.71, 0.55, respectively).

**Figure 1:**
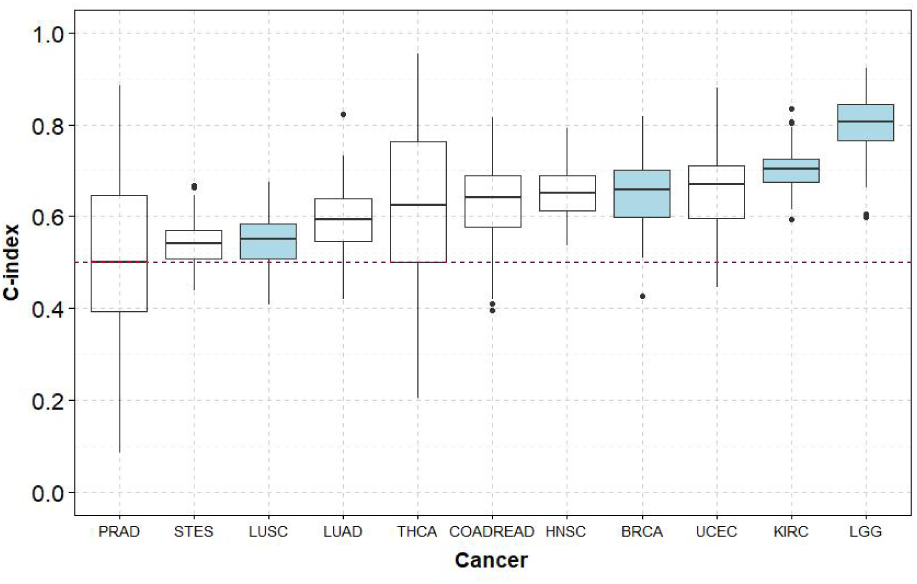
Boxplots of the C-indices for the 11 cancers from the TCGA with more than 500 patients with mRNA-seq data available, computed with the ridge penalty and all of the genes in real datasets. PRAD, prostate adenocarcinoma; STES, stomach and esophageal carcinoma; LUSC, lung squamous cell carcinoma; LUAD, lung adenocarcinoma; THCA, thyroid carcinoma; COADREAD, colorectal adenocarcinoma; HNSC, head and neck squamous cell carcinoma; BRCA, breast invasive carcinoma; UCEC, uterine corpus endometrial carcinoma; KIRC, kidney renal clear cell carcinoma; LGG, brain lower grade glioma.

The clinical and mRNA-seq datasets were obtained using the Broad GDAC FIREHOSE utility (https://gdac.broadinstitute.org/). We used the RNA-seq data normalized according to expectation maximization (RSEM) [20], and for all of the estimations of 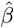 with the Cox regression models, the gene expression data were standardized. The Figures are shown for KIRC here in the main text, and the Figures for BRCA, LGG and LUSC are available as Supplementary Information.

### 2.6 Simulation pipeline and selection accuracy for different IQR thresholds

We set up a simulation pipeline to assess the selection accuracy compared to the so-called ground truth, which is graphically explained in Supplementary Figure S1 (R script *functions/simulations func.R*). First, we defined the ground truth as all of the genes selected in common by the lasso, elastic net, and adaptive elastic net, plus the genes selected by at least one of the three methods for the real TCGA dataset. We set the number *N* of genes of the ground truth as the median number of genes selected by the three methods, which corresponds to the number of genes selected by the elastic net. As an example, for KIRC and without pre-filtering, *N* = 55.

Characteristics of the four cancers used. C-indices were computed with the ridge regression and all of the genes in the Cox model, and *p*_3_ is the number of genes with an IQR *≥* 3.

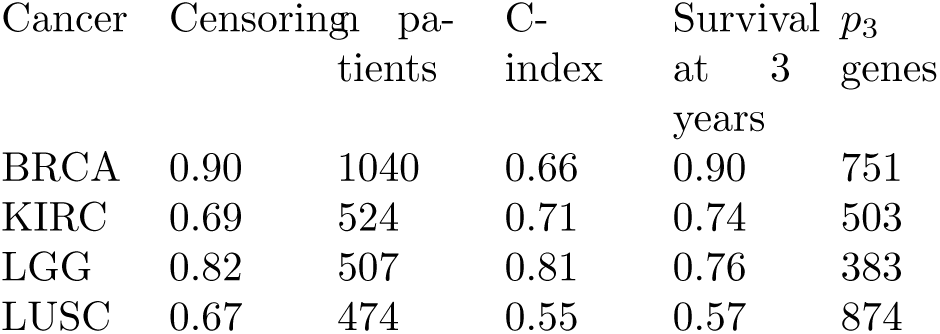

To obtain realistic simulations, we calibrated a Weibull proportional hazard model on real TCGA data using the genes of the ground truth. Then we simulated survival times *T*_*i*_, for *i* = 1, *…, n*, from TCGA mRNA-seq data using the [2] method. Censoring times *C*_*i*_, for *i* = 1, *…, n*, are assumed to be uniform, and the censoring rate of the real survival data is achieved using the method proposed by [37]. We define the follow-up times *t*_*i*_ as *t*_*i*_ = min(*T*_*i*_, *C*_*i*_), with *T*_*i*_ the survival time and *C*_*i*_ the censoring time, and the status 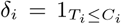, for *i* = 1, *…, n*. In the following, we will indicate whether the results are obtained from real or simulated datasets.

We estimate the selection accuracy of the four selection methods (i.e., lasso, elastic net, adaptive elastic net, univariate Cox) by comparing the set of selected genes for the simulated data with the set of genes of the ground truth, using the sensitivity and the false discovery proportion (FDP) (R script *selection_accuracy.R*). We were careful to select in the ground truth only genes that satisfied the chosen IQR threshold. As a result, the procedure naturally favors high IQR thresholds, as the selection is performed on fewer genes.

The genes selected by univariate Cox selection are defined as the *N* genes with the lowest p-values, where *N* is the number of genes in the ground truth. This scenario is then the most favorable for selection accuracy.

### 2.7 Stability of the prognostic index and reliability of the selected set of genes

We define a procedure as unstable if small changes in the input (patients) can cause large changes in the output (PIs, or set of genes with nonzero coefficients, also referred to as the ‘selected genes’).

To assess the robustness of the PIs, we compute 100 PIs for real TCGA datasets for each patient, using the bootstrapping method (i.e., random sampling with replacement). This makes it possible to compute an IQR and a median for each PI from their empirical bootstrap distribution. Then we fitted the loess curves to estimate the relation between IQR and the median PI. We repeat this procedure for each penalization method (i.e., lasso, elastic net, adaptive elastic net, ridge) (R script *stability_PI.R*).

For the stability of the selection, we randomly select genes from two separated sub-datasets of the real TCGA dataset that share *x* % common patients, with *x* ∈ {100, 90, 80, 70, 60, 50} while the sample size of patients remains constant. Then we compare the proportions of common genes selected from these two subdatasets. This procedure is graphically explained in Supplementary Figure S2. To further assess the reproducibility of the set of selected genes, the dataset is then divided into two subdatasets with no common patients (*x* = 0). Each procedure is repeated for 100 times (i.e, 100 Monte-Carlo runs), and the proportions of common genes are drawn as a function of the proportion of common patients for each selection method (i.e., lasso, elastic net, adaptive elastic net, univariate Cox) (R script *stability genes.R*).

### 2.8 Impact of the number of patients

To assess the impact of the number of patients in a cohort to infer 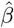, we use real TCGA datasets and we consider only a subset of *n*_*T*_ patients for the training set (*n*_*T*_ ∈ {25, 50, 100, 200, 400}), while keeping 100 patients in the test set to compute the C-index estimates (R script *impact_n_ patients.R*).

## 3 Results

### 3.1 Prediction quality is cancer dependent

Figure 1 shows the distribution of the C-index estimates for the 11 cancers using the estimation procedure described in section 2.4 for the real TCGA datasets. Prostate adenocarcinoma (PRAD) has the lowest prediction quality, with a median C-index of 0.50, which corresponds to pure random estimation. Different characteristics are observed among the four cancers we selected further. LGG shows the highest median concordance (*C* = 0.81), KIRC ranks second (*C* = 0.70), BRCA has intermediate predictive accuracy (*C* = 0.66), and LUSC has low median concordance (*C* = 0.55). This indicates that the overall survival prediction quality using a Cox model is cancer dependent.

### 3.2 Pre-filtering to keep only the most variable genes preserves prediction performance while reducing the dimension

For all of the four cancers investigated, with the removal of the genes with IQR ≥ 4, the C-indices computed for the real TCGA datasets vary very slightly (Fig. 2; Supplementary Fig. S3). However, filtering of more variable genes (IQR ≥ 5) leads to accuracy reductions in three of the four cancers investigated. For KIRC, this pre-filtering step allows a reduction from 20,245 to 178 for the number of genes, while hardly affecting the prediction accuracy. The median C-index even increases for BRCA: for the ridge regression, this goes from 0.66 without pre-filtering (20,248 genes) to 0.70 when restricted to the most variable genes across the patients (IQR ≥ 4, 271 genes).

**Figure 2:**
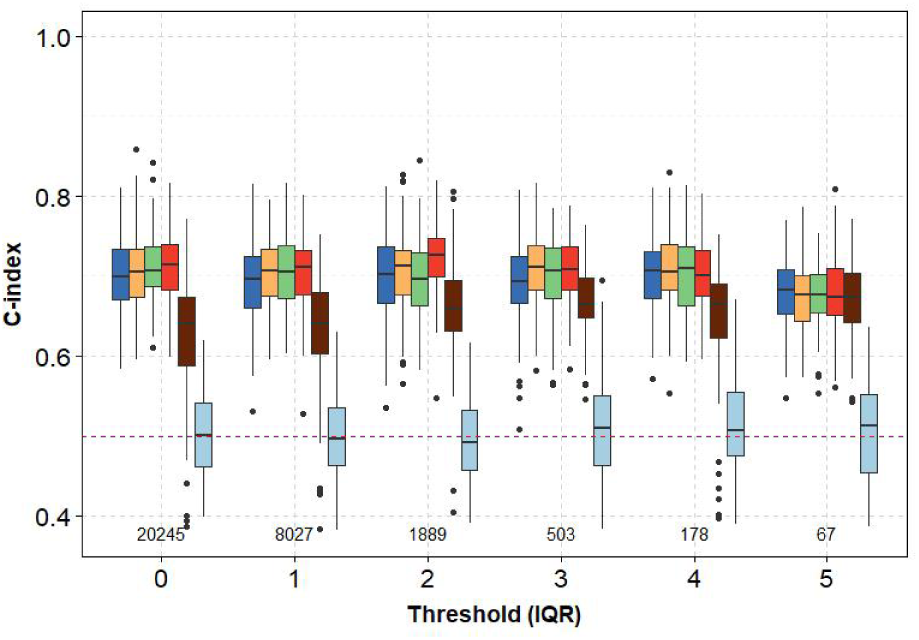
Prediction performance (C-index) for the real TCGA dataset for kidney renal clear cell carcinoma. Boxplots of C-indices as a function of IQR threshold. The numbers of genes selected by the pre-filtering step are also shown. Blue, the lasso; orange, the elastic net; green, the adaptive elastic; red, the ridge; dark red, the univariate Cox; light blue, the ridge under the null hypothesis.

We suggest that the C-index is used to determine the IQR thresholds for the pre-filtering step as follows:

- Determine the different thresholds to test according to the distribution of IQR for each gene (in this study, typically 0,1,…,5).
- Compute 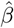 with the chosen method from a random selection of 80% of the patients, and estimate the C-index with the remaining 20% of the patients, 100 times for each threshold.
- Choose a threshold for the pre-filtering step according to the observed relation C=f(IQR threshold), as in Figure 2.

Another important feature is that the elastic net is less stringent than the lasso, as it selects more genes, while it is more stringent than the adaptive elastic net (Supplementary Fig. S4). For the LUSC patients analyzed with the lasso and elastic net penalties, keeping all of the genes leads to no genes selected in many cases (Supplementary Fig. S4D). For the adaptive elastic net, due to the weight on each gene that is calculated in the first step of the procedure, a few tens of genes are always selected, while the overall performance remains very low, with a concordance of ∼ 0.50. For this cancer, the pre-filtering of the genes clearly improves the estimation, but the C-index remains typically below 0.55, with an optimal threshold of 4 or 5, depending on the penalization method (Supplementary Fig. S3C).

For all of the IQR thresholds, the median C-indices are significantly higher for all multivariate methods compared to the univariate Cox selection for KIRC (Fig. 2) and LGG (Supplementary Fig. S3B), and equivalent for BRCA and LUSC (Supplementary Figs. S3A, S3C). A linear combination of gene signatures with coefficients estimated in the Cox model provides better prediction of the overall survival than one gene alone. The different multivariate Cox penalizations (i.e., lasso, elastic net, adaptive elastic net, ridge) perform equally in terms of prediction, but the lasso, elastic net and adaptive elastic net penalties allow for the selection of models of moderate size.

The C-indices computed under the null hypothesis “*β* = 0” for each methodology and each threshold are shown in Supplementary Figure S5 for KIRC, and these allow comparisons of the results with the pure random model.

### 3.3 Gene selection leads to high rate of false positives and negatives and low stability performance

We assess the selection performance using simulations of overall survival and follow-up time. Best results are obtained with the genes with an IQR ≥ 4.The ground truth, as the pool of genes used for this simulation, is comprised of 37 genes for KIRC that are chosen from among the 178 genes with IQR ≥ 4, as detailed in the Materials and Methods section. The elastic net reaches the highest median sensitivity (0.70), but with a median FDP of 0.55 (Fig. 3). The lasso and adaptive elastic net penalties perform similarly, with sensitivities of 0.58 and 0.62, respectively, and FDPs of 0.48 and 0.49, respectively. Indeed, overall, half of the selected genes are false positives, and 75% of the ground truth is discovered in the best scenario. The univariate Cox models clearly perform the worst to retrieve the genes of the ground truth, with a sensitivity below 25% and an FDP above 75%. Similar results are obtained for BRCA, LGG, and LUSC (Supplementary Fig. S6). Supplementary Figure S7 shows that the elastic net and adaptive elastic net penalties select too many genes, while the lasso selects a number of genes comparable to the length of the ground truth.

**Figure 3:**
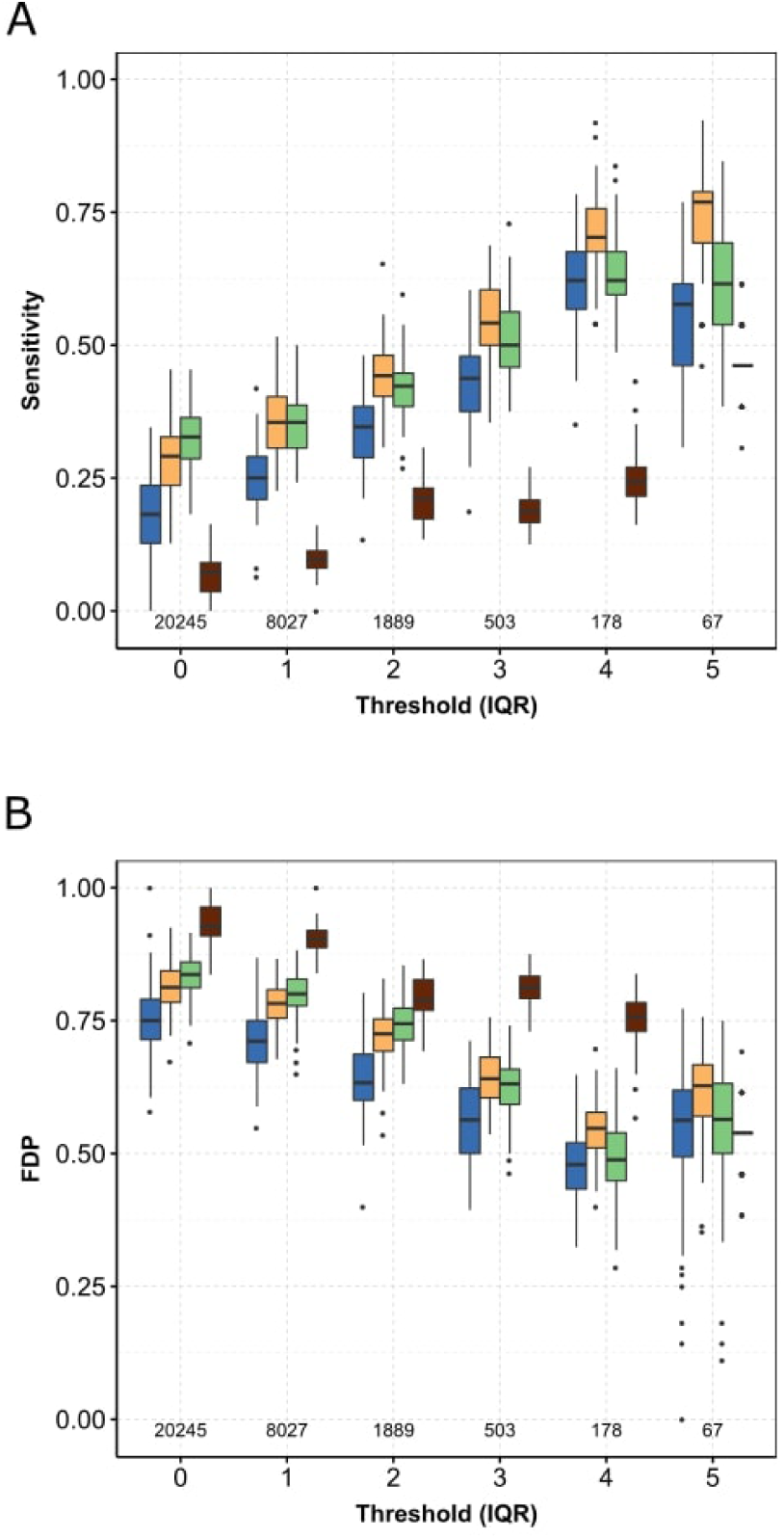
Selection accuracy for the simulated dataset estimated for kidney renal clear cell carcinoma. Boxplots of sensitivity (A), false discovery proportion (FDP) (B), computed on simulated datasets as a function of the IQR threshold. Blue, the lasso; orange, the elastic net; green, the adaptive elastic net; dark red, univariate Cox.

A good compromise in terms of concordance and selection performance among the five methods (i.e., lasso, elastic net, adaptive elastic net, ridge, univariate Cox) and the four studied cancers is a threshold of 3 for the IQR. This reduces the number of genes to 300 to 900, depending on the cancer. For the following, we use a pre-filtering step, to only keep the genes with IQR ≥ 3.

### 3.4 The ridge regression reduces the PIs more and is more robust than the lasso, elastic net and adaptive elastic net penalties

Figure 4 shows that the PIs computed with the ridge penalty for the real TCGA subdatasets are more robust and reduced than for the three other methods (i.e., lasso, elastic net, adaptive elastic net). Among the three selection methods, the PIs computed with the elastic net have the best robustness, while the adaptive elastic net performs the worst. Similar results are obtained for BRCA, LGG and LUSC (Supplementary Fig. 8). The squared *ℓ*_2_-norm in the elastic net penalty brings stability to the estimation of 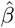, whereas the two steps of the adaptive elastic net makes the estimation more unstable.

**Figure 4:**
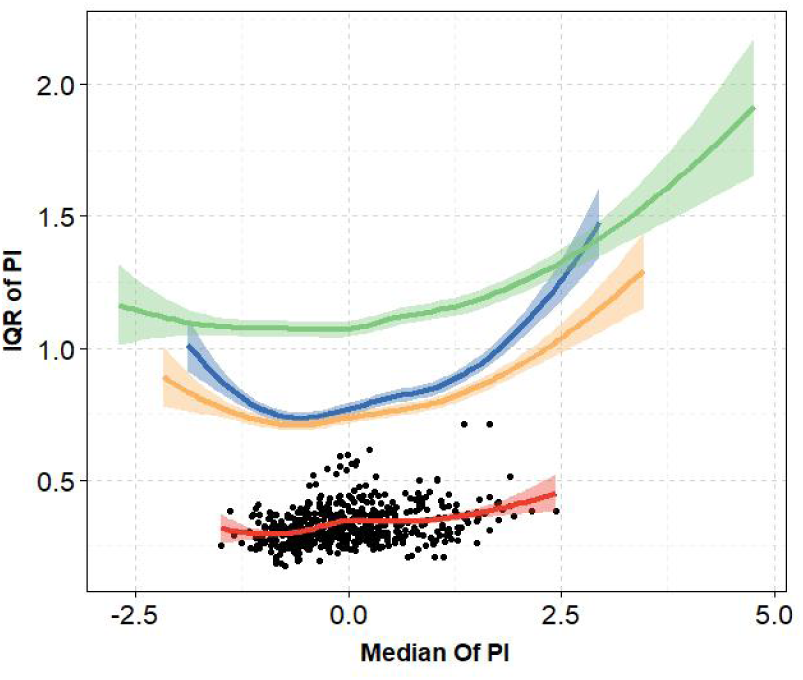
Robustness of the Prognostic Indices (PIs) estimated with the genes with IQR *≥* 3 for the real TCGA dataset for kidney renal clear cell carcinoma. Loess curves for the IQR of the PIs as a function of the median PIs. The black symbols are associated with the red curve. The PIs are estimated by cross-validation with the ridge and bootstrap. Blue, the lasso; orange, the elastic net; green, the adaptive elastic; red, the ridge.

### 3.5 Gene selection is unstable

To evaluate the stability of the pool of selected genes using the different variants of the lasso, we compare the pools obtained for two subsets of the real TCGA data. A decrease in the proportion of common patients quickly reduces the proportion of common genes selected, down to 25% for 50% of the patients in common (Fig. 5). When selecting genes within two subsets with independent patients, the proportion of overlapping genes drops to ¡10% for the penalized Cox models (6.7% for the lasso, 9.2% for the elastic net, 9.3% for the adaptive elastic net). Similar results are obtained for BRCA and LUSC. The best results remain ¡25%, and are obtained for LGG (13.8% for the lasso, 23.5% for the elastic net, 19.1% for the adaptive elastic net) (Supplementary Fig. S9). The univariate Cox model reaches a higher stability performance, but does not exceed 50% of common genes selected. Note that for identical datasets (100% of common patients), the pools of genes selected are not exactly the same, as the inferred *λ*_*min*_ can differ due to the random partitions used in the cross-validation step.

**Figure 5:**
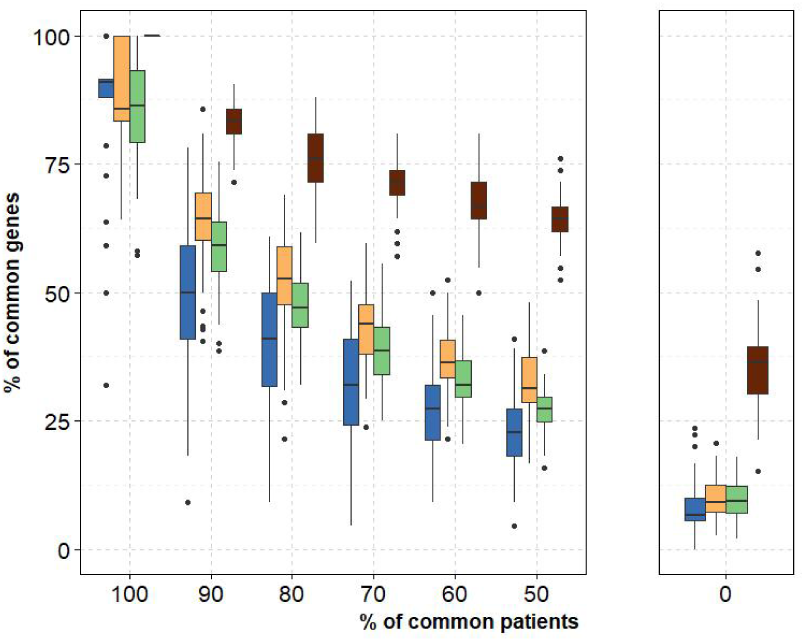
Stability of the genes selected computed with the genes with IQR *≥* 3 from the real TCGA dataset for kidney renal clear cell carcinoma. Boxplots of proportions of common genes selected in the two datasets as a function of the proportion of common patients in the two datasets. Blue, the lasso; orange, the elastic net; green, the adaptive elastic net; dark red, the univariate Cox.

### 3.6 Prediction accuracy reaches a plateau from different numbers of patients, which depends on the cancer

As expected, concordance increases with the number of patients used to estimate 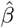 for the real TCGA datasets (Supplementary Fig. S10). This reaches a plateau in different ways, depending on the cancer considered. For KIRC, the plateau is not even clearly reached with 400 patients, whereas for LGG the plateau is reached for 100-200 patients. Except for LGG, for which the ridge regression performs similarly with the other penalized Cox methods, the ridge method slightly outperforms the other penalization methods when the numbers of patients are low. As the cover is different for each *n*_*T*_ (i.e., between each bootstrap runs, there are more common patients in the training sets for *n*_*T*_ = 400 than for *n*_*T*_ = 25), the variances are biased. Thus, we only show the median C-indices here.

## 4 Discussion

We first show here that overall survival prediction quality is cancer dependent, and can be as low as pure random prediction, with a median concordance of 0.50 for PRAD. Also, for the univariate selection procedures based on correlation between covariates and binary outcome (”good” or “bad”), the prognostic genes listed were shown to be non-robust [10]. In a similar way, our results illustrate the same issue in multivariate Cox selection methods. Different hypotheses can explain both these differences across cancers and the poor selection performances. First, many studies show intra-tumoral heterogeneity (e.g., genetic and phenotypic variations across different geographic regions for one tumor), including for brain, breast, renal, and lung cancers [33]. The expression levels of the genes correlated to survival (the input of the Cox model) can then vary across geographic regions for one tumor, although the survival times of the patients are a global outcome. Secondly, other explanatory variables can better predict overall survival for some cancers. For example, [30] recently showed that tumor microbiome diversity influences the tumor outcome for patients with pancreatic cancer. Predicting patient outcome with transcriptomic data if they are poorly correlated to these hidden variables here means a low probability of success. Finally, the existence of sub-types of cancers with different molecular characteristics [7, 19] would lead to poor regression performance and prediction, as all of the patient data are treated similarly with the same model.

The main goal of penalty methods in Cox models is both to provide significant weight for prognostic genes, and to predict overall survival. Several comparisons of selection methods that include multivariate Cox models have been carried out, but these were applied to smaller datasets and with different metrics to assess the quality of the models. [3] concluded that there is a strong need for research on improved pre-filtering of covariates for high-dimensional data Cox regression. We share these conclusions, and we propose a methodology to tune the pre-filtering of the genes based on a robust estimation of their variability among biopsies from patients.

Even if direct application of the lasso alternatives of the Cox model shows disappointing prediction for many cancer types, promising strategies do show up. [8] suggested to implement ‘knowledge-based’ (as opposed to ‘ignorance-based’) approaches, as widely used in all branches of physics. Bioinformatics associated with strong clinical and biological assumptions can increase the robustness, simplicity, and accuracy of such results. Finally, the author also showed that combining ‘omics’ and clinical data showed promising results.

## 5 Conclusions

This article investigates Cox model penalty methods (i.e., lasso, elastic net, adaptive elastic net, ridge), univariate Cox procedure, and pre-filtering of the genes based on their variability, for prediction of overall survival and the selection of a set of prognostic biomarkers. We compare four cancers (i.e., KIRC, BRCA, LGG, LUSC) using data from TCGA. We first demonstrate that the prediction accuracy is highly cancer dependent, with concordance indices as high as 0.81 for LGG, to 0.55 for LUSC, and down to 0.50 for PRAD. Secondly, we propose a methodology to pre-filter the genes according to their variability, and show that the prediction accuracy varies very slightly while keeping the genes with IQR ≥ 4. This methodology allows for an important dimension reduction. Thirdly, we show that even in the ideal situation with the data simulated using the Cox model, the pool of selected genes is not stable, with many false positive and false negative genes. Fourth, multivariate Cox penalties (i.e., lasso, elastic net, adaptive elastic net, ridge) perform better than the univariate Cox procedure for prediction of patient outcome, and the lasso, elastic net and adaptive elastic net penalties are also more powerful for selection accuracy than the univariate Cox selection. Finally, we demonstrate that prognostic indices computed with the ridge regression are more robust than those computed with the lasso, elastic net or adaptive elastic net penalties, and they reach a slightly better prediction performances. Thus, we can be more confident with the use of the ridge penalty for computing the risk factor (PIs).

As practical conclusions, we first advise that the genes are pre-filtered based on their variability across patients, with the C-index computed by cross-validation as a decision criterion. Then, we do not advise to use selection procedures based on the Cox model to identify prognostic biomarkers. Finally, we advise to use the ridge penalty to predict overall survival after the pre-filtering step, and elastic net if a more parsimonious model is needed although the Prognostic Indices are less robust.

## Supporting information

Supplementary materials and R script

## Acknowledgements

The authors wish to thank Christopher Berrie for language editing.

## References

[1] Y. S. Abu-Mostafa et al., Learning From Data, Wiley Series in Probability and Statistics, 2012.

[2] R. Bender et al., Generating survival times to simulate Cox proportional hazards modelse, Statistics in Medicine, 24 (2005), pp. 1713–1723.

[3] A. Benner et al., High-dimensional cox models: The choice of penalty as part of the model building process, Biometrical Journal, 52 (2010), pp. 50–69.

[4] R. Bourgon et al., Independent filtering increases detection power for high-throughput experiments, Proceedings of the National Academy of Sciences, 107 (2010), pp. 9546–9551.

[5] N. Breslow, Contribution to the Discussion of the Paper by D.R. Cox, Journal of the Royal Statistical Society B, 34 (1972), pp. 2016–2017.

[6] D. R. Cox, Regression Models and Life-Tables, Journal of the Royal Statistical Society. Series B: Statistical Methodology, 34 (1972), pp. 187–220.

[7] C. J. Creighton et al., Comprehensive molecular characterization of clear cell renal cell carcinoma, Nature, 499 (2013), pp. 43–49.

[8] E. Domany, Using high-throughput transcriptomic data for prognosis: A critical overview and perspectives, Cancer Research, 74 (2014), pp. 4612–4621.

[9] E. I. Dumbrava and F. Meric-Bernstam, Personalized cancer therapy — leveraging a knowledge base for clinical decision-making, Molecular Case Studies, 4(2) (2018), p. a001578.

[10] L. Ein-Dor et al., Outcome signature genes in breast cancer: is there a unique set?, Bioinformatics, 21 (2004), pp. 171–178.

[11] B. Fa and k.. others, Pathway-based biomarker identification with crosstalk analysis for robust prognosis prediction in hepatocellular carcinoma, EBioMedicine, 44 (2019), pp. 250 – 260.

[12] J. Friedman et al., Regularization paths for generalized linear models via coordinate descent, Journal of Statistical Software, 33 (2010), pp. 1–22.

[13] A. J. Hackstadt and A. M. Hess, Filtering for increased power for microarray data analysis, BMC Bioinformatics, 10 (2009), p. 11.

[14] F. E. Harrell Jr. et al., Multivariable prognostic models: Issues in developing models, evaluating assumptions and adequacy, and measuring and reducing errors, Statistics in Medicine, 15 (1996), pp. 361–387.

[15] L. Hood and S. H. Friend, Predictive, personalized, preventive, participatory (P4) cancer medicine, Nature Reviews Clinical Oncology, 8 (2011), pp. 184–187.

[16] R. Jardillier et al., Bioinformatics Methods to Select Prognostic Biomarker Genes from Large Scale Datasets : A Review, Biotechnology Journal, 13 (2018), pp. 1–12.

[17] J. D. Kalbfleisch and R. L. Prentice, The Statistical Analysis of Failure Time Data, AMLBook, 2011.

[18] M. W. Kattan, Comparison of cox regression with other methods for determining prediction models and nomograms, The Journal of Urology, 170 (2003), pp. S6 – S10. Part 2 of 2.

[19] D. C. Koboldt et al., Comprehensive molecular portraits of human breast tumours, Nature, 490 (2012), pp. 61–70.

[20] B. Li and C. N. Dewey, Rsem: accurate transcript quantification from rna-seq data with or without a reference genome, BMC Bioinformatics, 12 (2011), p. 323.

[21] Q. Liao et al., Large-scale prediction of long non-coding RNA functions in a coding–non-coding gene co-expression network, Nucleic Acids Research, 39 (2011), pp. 3864–3878.

[22] S. Michiels et al., Prediction of cancer outcome with microarrays: a multiple random validation strategy, The Lancet, 365 (2005), pp. 488 – 492.

[23] L. D. Miller et al., Optimal gene expression analysis by microarrays, Cancer Cell, 2 (2002), pp. 353 – 361.

[24] F. M. Ojeda et al., Comparison of cox model methods in a low-dimensional setting with few events, Genomics, Proteomics Bioinformatics, 14 (2016), pp. 235 – 243.

[25] M. Pavlou et al., How to develop a more accurate risk prediction model when there are few events, BMJ, 351 (2015).

[26] M. J. Pencina and R. B. D’Agostino, Overall c as a measure of discrimination in survival analysis: model specific population value and confidence interval estimation, Statistics in Medicine, 23 (2004), pp. 2109–2123.

[27] R Core Team, R: A Language and Environment for Statistical Computing, R Foundation for Statistical Computing, Vienna, Austria, 2019.

[28] P. Raman et al., A comparison of survival analysis methods for cancer gene expression rna-sequencing data, Cancer Genetics, 235-236 (2019), pp. 1 – 12.

[29] S. Ramaswamy et al., A molecular signature of metastasis in primary solid tumors, Nature Genetics, 33 (2003), pp. 49–54.

[30] E. Riquelme et al., Tumor microbiome diversity and composition influence pancreatic cancer outcomes, Cell, 178 (2019), pp. 795 – 806.e12.

[31] M. Schroeder et al., survcomp: an r/bioconductor package for performance assessment and comparison of survival models, Bioinformatics, 27(22) (2011), pp. 3206–3208.

[32] T. Sorlie et al., Gene expression patterns of breast carcinomas distinguish tumor subclasses with clinical implications, Proceedings of the National Academy of Sciences of the United States of America, 98 (2001), pp. 10869–10874.

[33] X.-x. Sun and Q. Yu, Intra-tumor heterogeneity of cancer cells and its implications for cancer treatment, Acta Pharmacologica Sinica, 36 (2015), pp. 1219–1227.

[34] K. Suresh and S. Chandrashekara, Sample size estimation and power analysis for clinical research studies, Journal of Human Reproductive Sciences, 5 (2012), pp. 7–13.

[35] R. Tibshirani, The lasso method for variable selection in the cox model, Statistics in Medicine, 16 (1997), pp. 385–395.

[36] P. J. M. Verweij and H. C. Van Houwelingen, Penalized likelihood in cox regression, Statistics in Medicine, 13 (1994), pp. 2427–2436.

[37] F. Wan, Simulating survival data with predefined censoring rates for proportional hazards models, Statistics in Medicine, 36 (2017), pp. 838–854.

[38] A. Xiang et al., Comparison of the performance of neural network methods and cox regression for censored survival data, Computational Statistics Data Analysis, 34 (2000), pp. 243 – 257.

[39] H. Zou, The adaptive lasso and its oracle properties, Journal of the American Statistical Association, 101 (2006), pp. 1418–1429.

[40] H. Zou and T. Hastie, Regularization and variable selection via the elastic-net, Journal of the Royal Statistical Society, 67 (2005), pp. 301–320.

[41] H. Zou and H. H. Zhang, On the adaptive elastic-net with a diverging number of parameters, Annals of Statistics, 37 (2009), pp. 1733–1751.

