## Supplementary materials and R script for "Benchmark of lasso-like penalties in the Cox model for TCGA datasets reveal improved performance with pre-filtering and wide differences between cancers": READ_ME.txt

This folder contains:

-"code.zip": the code used to generate the figures.

-"main_figures_high_resolution.zip": low resolution figures are included within the text, and high resolution 
figures are available in this folder.

- "Supplementary_figures.pdf": Supplementary figures with titles.
