## Supplementary materials and R script for "Benchmark of lasso-like penalties in the Cox model for TCGA datasets reveal improved performance with pre-filtering and wide differences between cancers": Supplementary_figures.pdf

**A**

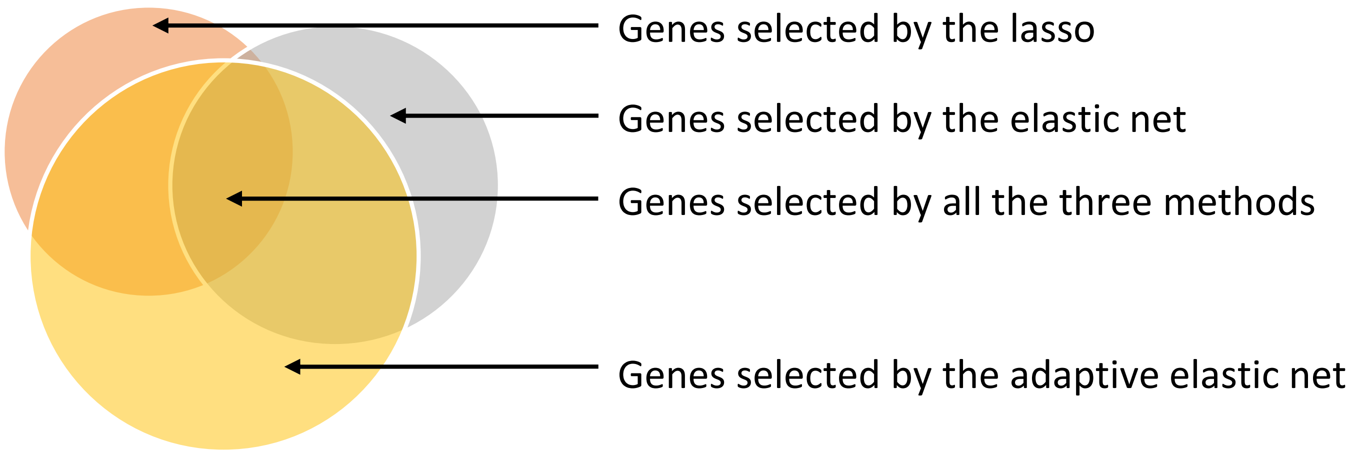


(1) Ground truth: genes selected by all three of the methods + genes selected by at least one of the methods.
(2) Card (ground truth): Card (genes selected by the elastic net), with # as the function that returns the number of elements in a set.

**B**

(1) Weibull-Cox model calibrated on real TCGA (*coxph* function from *survival* package) dataset with the genes from the ground truth: $h(t) = \lambda p t^{p-1} \exp(\beta \boldsymbol{X})$
$\beta_i = 0$ if the gene i is not in the ground truth
$\beta_i \neq 0$ if the gene i is in the ground truth

(2) $\hat{\beta}$, λ, p, X, —*Bender et al., 2005*→ $T^{(j)} = \left(-\frac{\log(U)}{\lambda \exp(\beta^T \boldsymbol{X}^{(j)})}\right)^{1/p}$, the overall survival for patient j, with U a random number between 0 and 1.

With X the gene expression matrix from TCGA

↓ *Wan, 2016*

(3)

$$C_j \sim U[0, \theta]$$
$$\hat{\theta} = \arg\min_{\theta} \left(\frac{1}{n} \sum_{i=j}^{n} P(C_j < T_j | \boldsymbol{X}^{(j)}, \theta, \lambda, k) - c_r\right)^2$$
$$P(C_j < T_j | \boldsymbol{X}^{(j)}, \theta, \lambda, k) = \frac{1}{p\theta\lambda^{1/p} \exp(\beta^T \boldsymbol{X}^{(j)})^{1/p}} \gamma\left(\frac{1}{p}, \lambda\theta^p \exp(\beta^T \boldsymbol{X}^{(j)})\right)$$
$$\delta_j = 1_{T_j < C_j}$$

**Supplementary Fig. 1. Simulation pipeline.** (A) Construction of the ground truth with the genes selected by the lasso, the elastic net and the adaptive elastic net. (B) The three steps of the simulation pipeline: (1) Calibration of a Weibull-Cox model on the real TCGA dataset with genes from the ground truth; (2) Simulations of overall survival with the Bender et al. method for all patients; (3) Simulations of the censoring times with the Wan method for all patients, and definition of status.

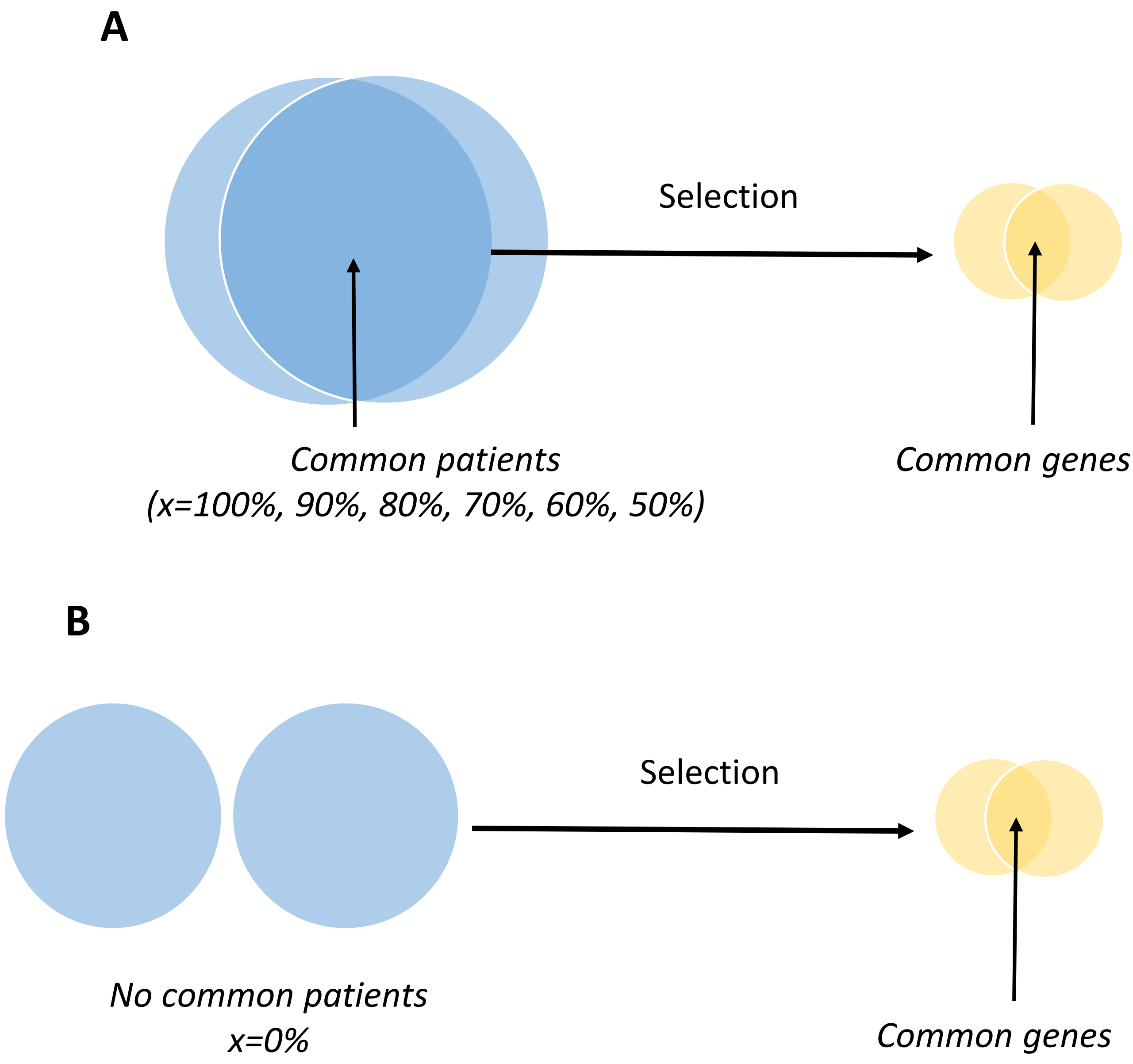


**Supplementary Fig. 2. Methodology to estimate the stability of the genes selected.** (A) Two subdatasets sharing x% common patients (blue), and genes selected in common for these two subdatasets (yellow). (B) Two subdatasets that do not have common patients (blue), and genes selected in common for these two subdatasets (yellow).

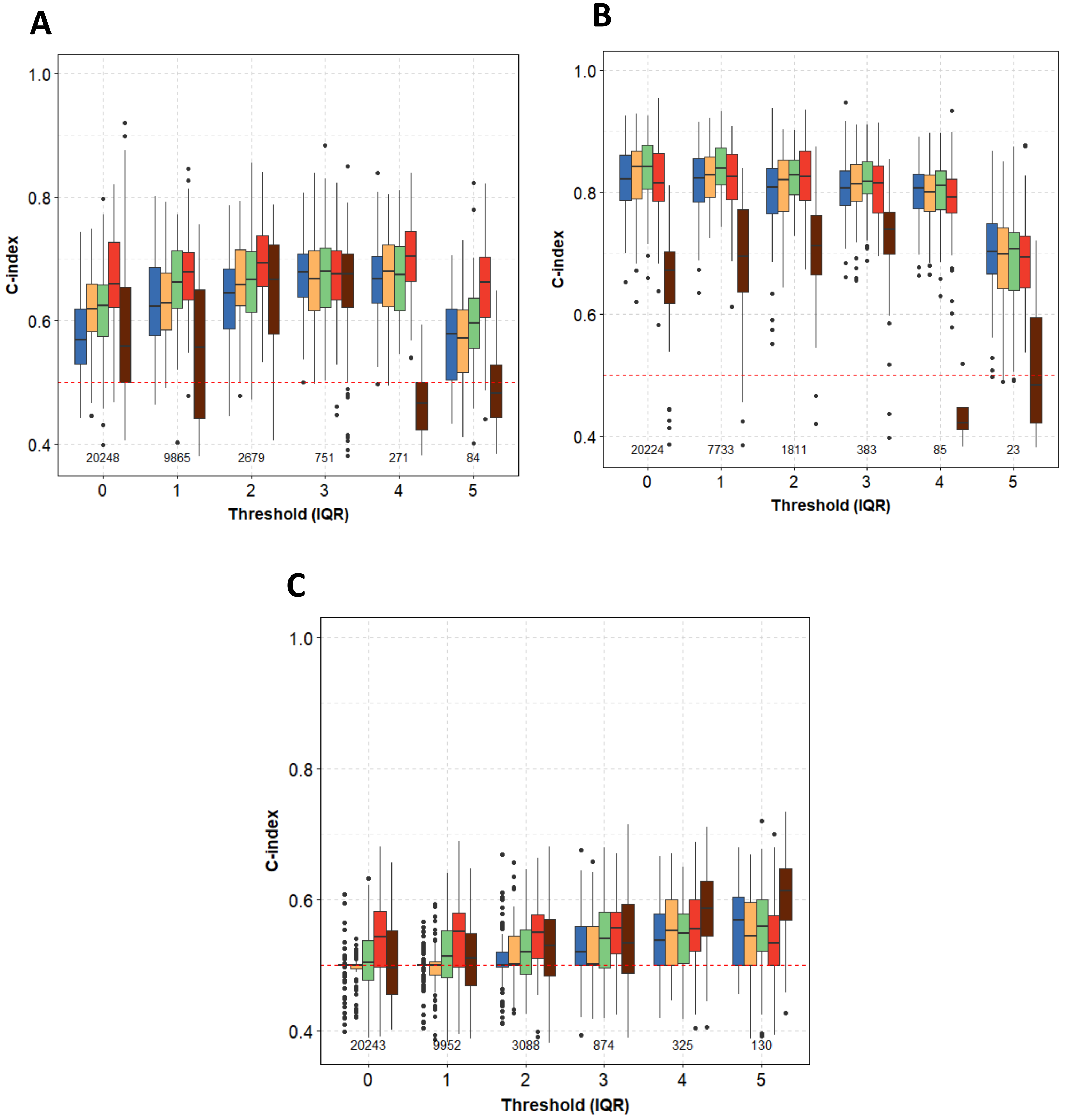


**Supplementary Fig. 3. Prediction performance (C-index) for the real TCGA dataset for breast invasive carcinoma (A), brain lower grade glioma (B), and lung squamous cell carcinoma (C).** Boxplots of C-indices as a function of IQR threshold. The numbers of genes selected by the pre-filtering step are also shown. Blue, the lasso; orange, the elastic net; green, the adaptive elastic; red, the ridge; dark red, the univariate Cox.

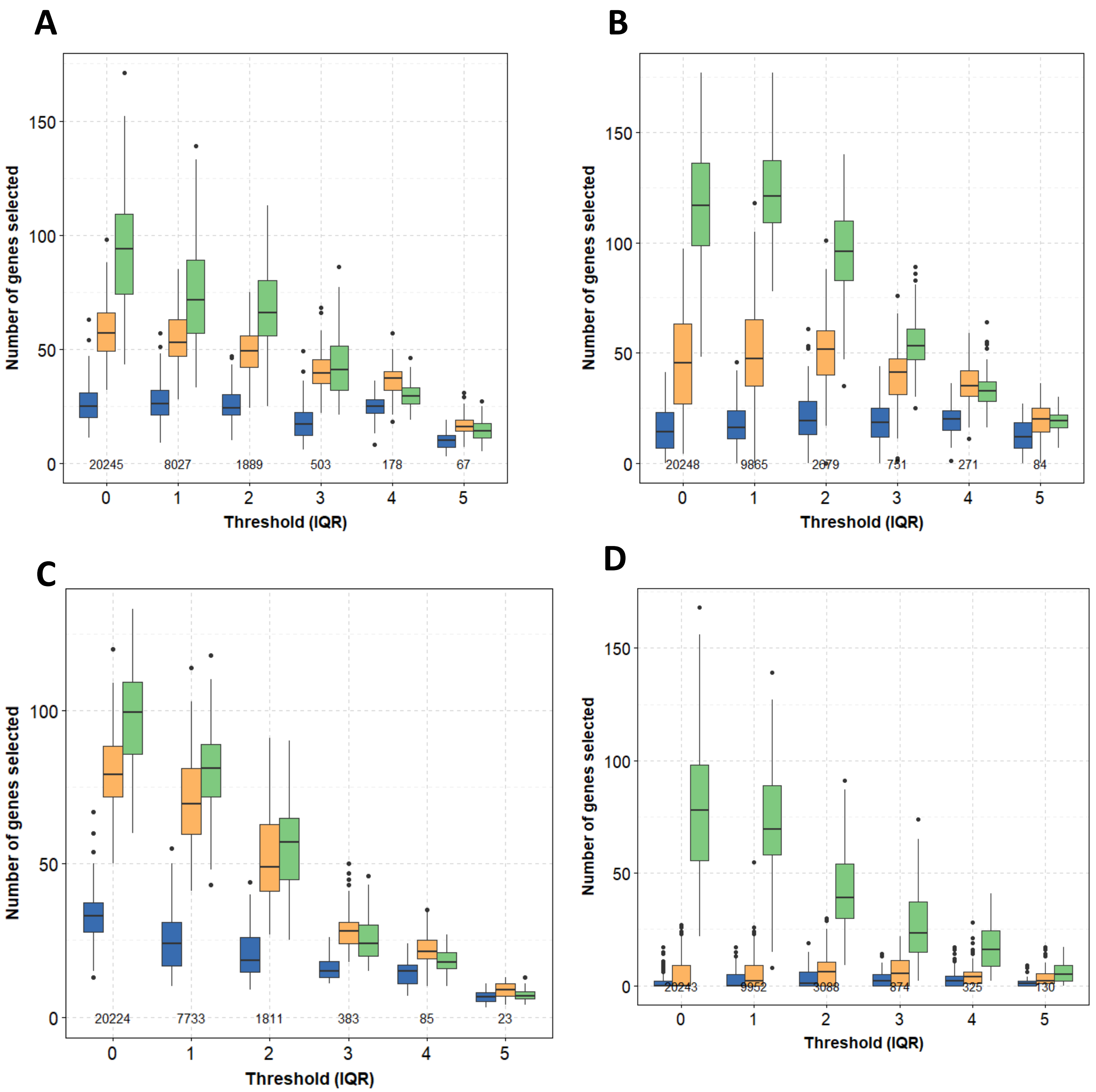


**Supplementary Fig. 4. Numbers of genes selected for the real TCGA dataset for kidney renal clear cell carcinoma (A), breast invasive carcinoma (B), brain lower grade glioma (C), and lung squamous cell carcinoma (D).** Boxplots of the numbers of genes selected computed on 80% of the patients as a function of the IQR threshold, for the lasso (blue), the elastic net (orange), the adaptive elastic net (green), the ridge (red), and the univariate Cox (dark red). The numbers of genes selected in the pre-filtering step are also shown.

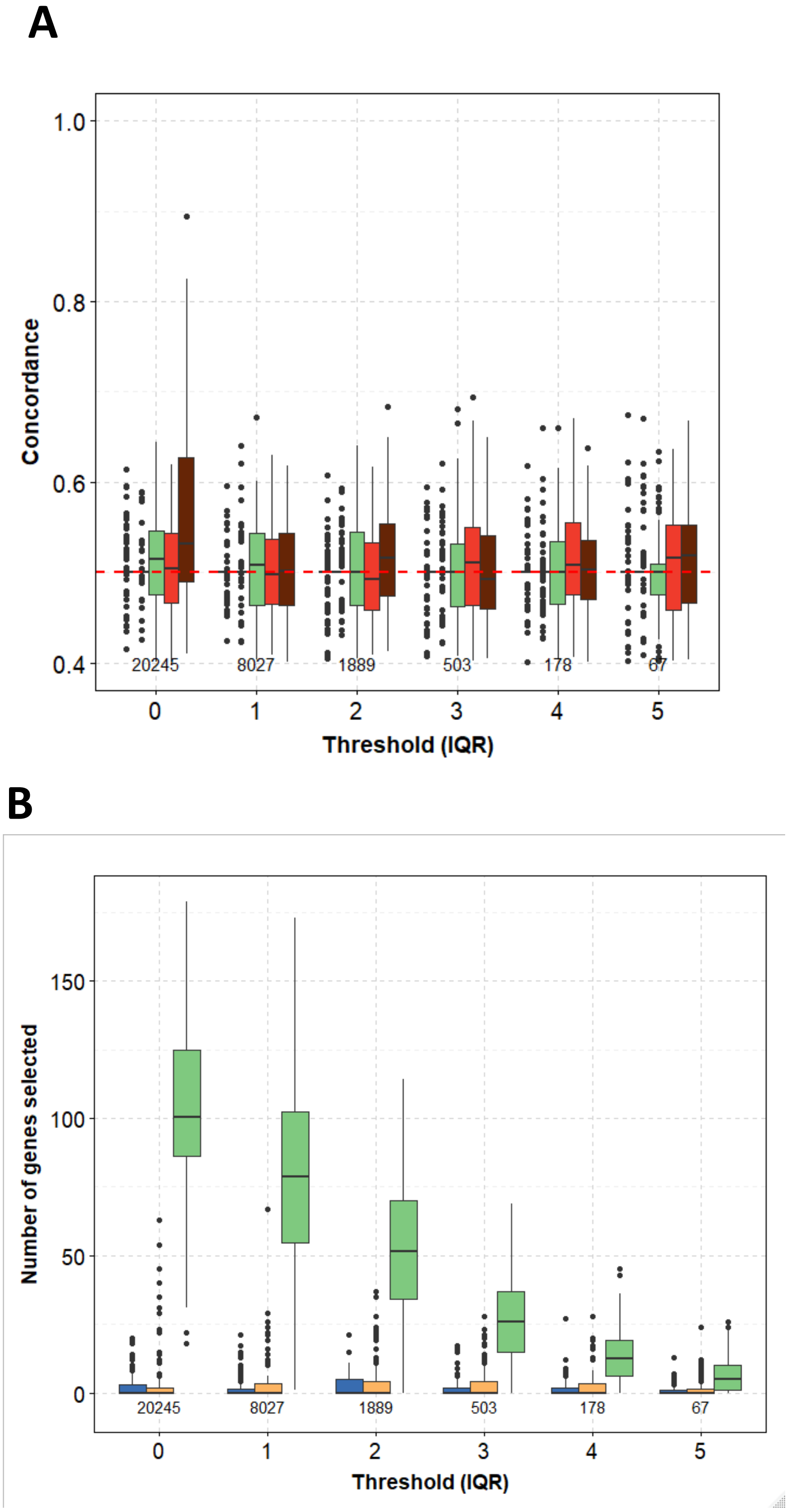


**Supplementary Fig. 5. Prediction performance (C-index) for the real TCGA dataset for kidney renal clear cell carcinoma under the null hypothesis ("There is no link between overall survival and mRNA expression") .** (A) Boxplots of the C-indices under the null hypothesis as a function of the IQR threshold. (B) Number of genes selected under the null hypothesis computed on 80% of the patients as a function of the IQR threshold. Blue, the lasso; orange, the elastic net; green, the adaptive elastic net; red, the ridge; dark red, the univariate Cox. The numbers of genes selected in the pre-filtering step are also shown.

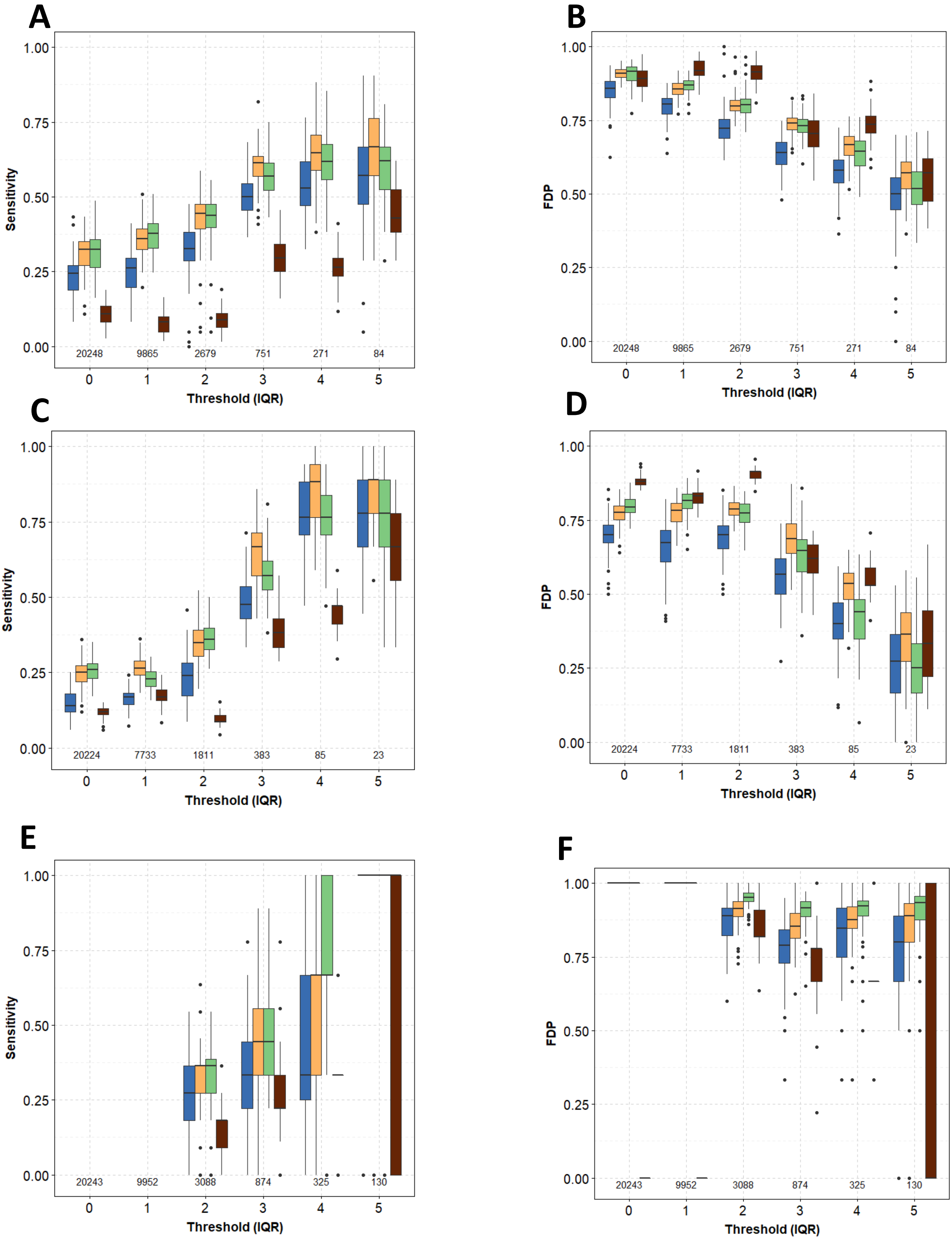


**Supplementary Fig. 6. Selection accuracy for the simulated dataset for breast invasive carcinoma (A, B), brain lower grade glioma (C, D), and lung squamous cell carcinoma (E, F).** Boxplots of sensitivity (A, C, E) and false discovery proportion (FDP) (B, D, F) as a function of the IQR threshold, for the lasso (blue), the elastic net (orange), the adaptive elastic net (green), and the univariate Cox (dark red). The numbers of genes selected in the pre-filtering step are also shown.

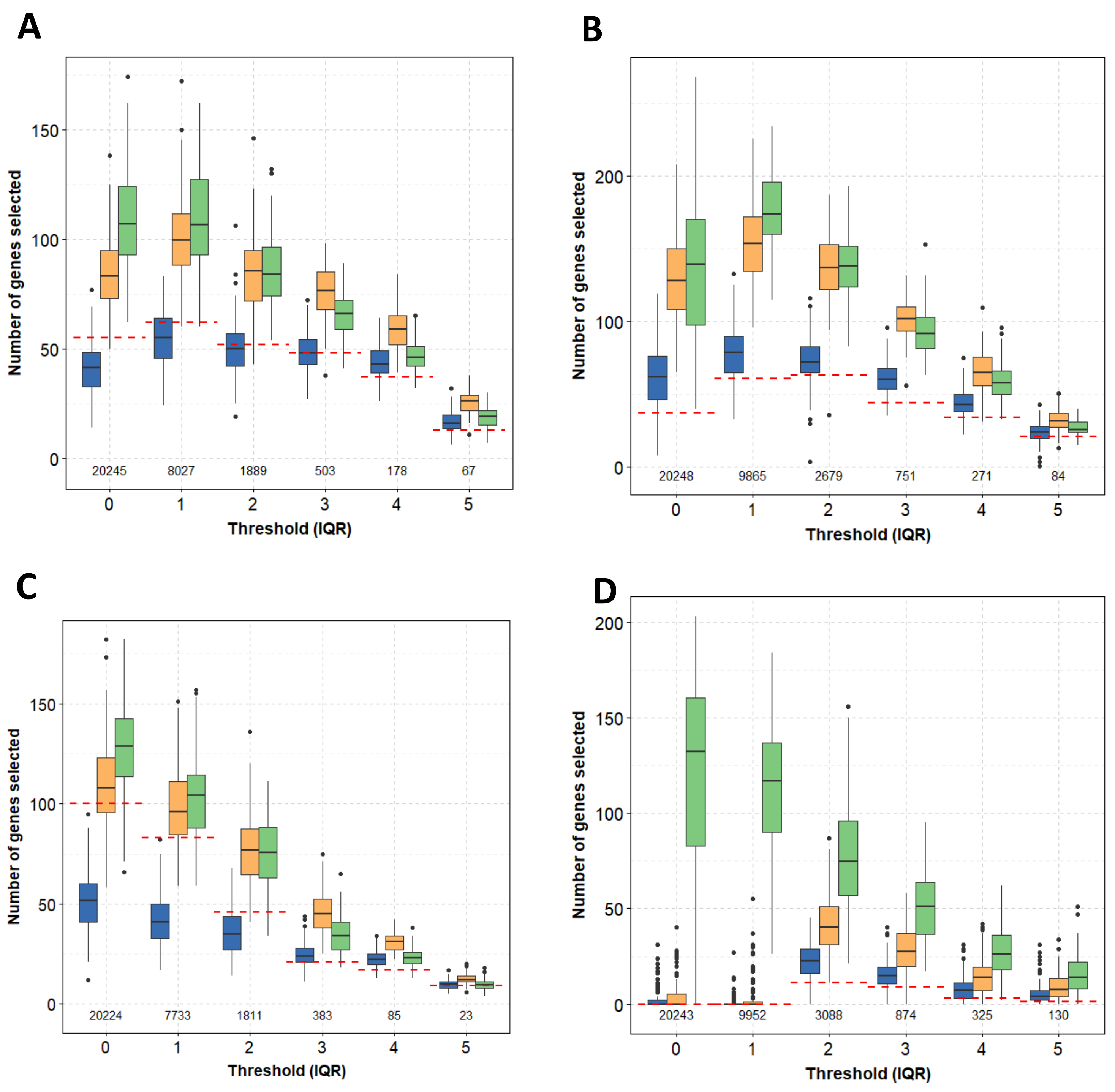


**Supplementary Fig. 7. Number of genes selected for the simulated dataset for kidney renal clear cell carcinoma (A), breast invasive carcinoma (B), brain lower grade glioma (C), and lung squamous cell carcinoma (D).** Boxplots of the numbers of genes selected in simulations as a function of the IQR threshold, for the lasso (blue), the elastic net (orange), and the adaptive elastic net (green). Red dotted lines, number of genes in the ground truth. The numbers of genes selected in the pre-filtering step are also shown.

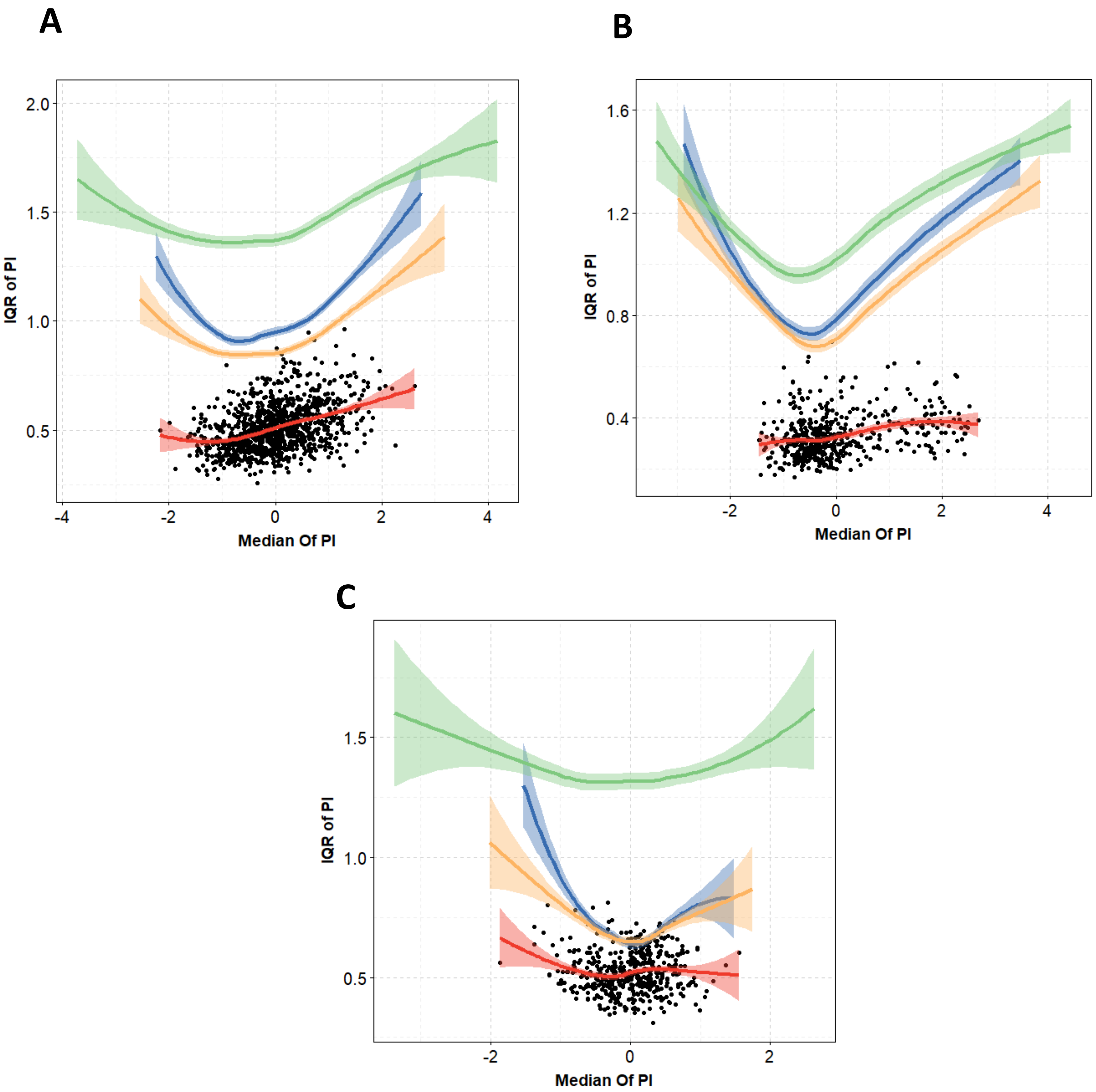


**Supplementary Fig. 8. Robustness of the Prognostic Indices (PIs) estimated with the genes with IQR ≥ 3 for the real TCGA dataset for breast invasive carcinoma (A), brain lower grade glioma (B), and lung squamous cell carcinoma (C).** Loess curves for the IQR of the PIs as a function of the median PIs. The black symbols are associated with the red curve. The PIs are estimated with the ridge and bootstrap. Blue, the lasso; orange, the elastic net; green, the adaptive elastic; red, the ridge.

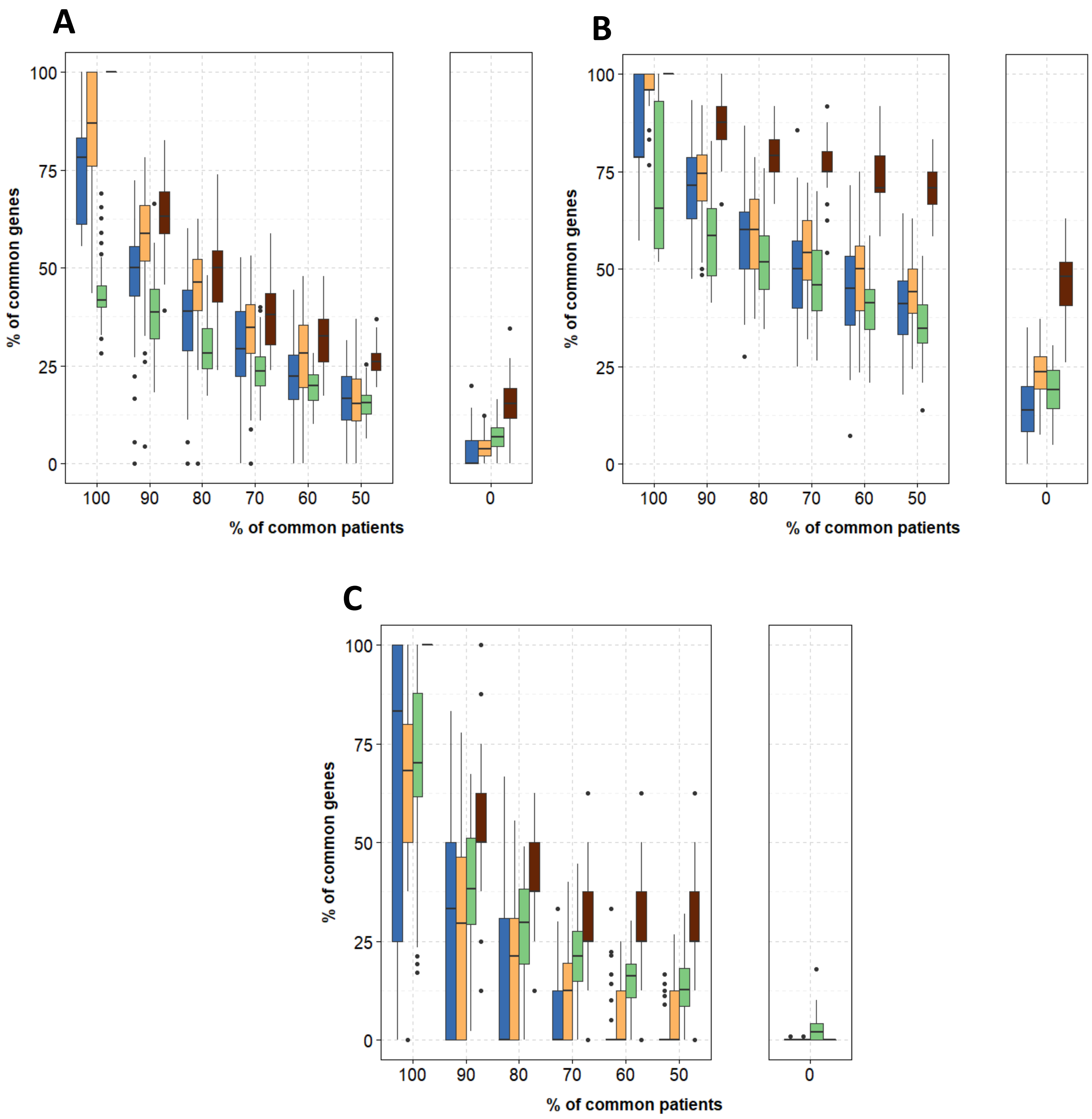


**Supplementary Fig. 9. Stability of the genes selected computed with the genes with IQR ≥ 3 for the real TCGA dataset for breast invasive carcinoma (A), brain lower grade glioma (B), and lung squamous cell carcinoma (C).** Boxplots of the proportions of the common genes selected in the two subdatasets as a function of the proportions of the common patients in these two subdatasets, for the lasso (blue), the elastic net (orange), the adaptive elastic net (green), and the univariate Cox (dark red).

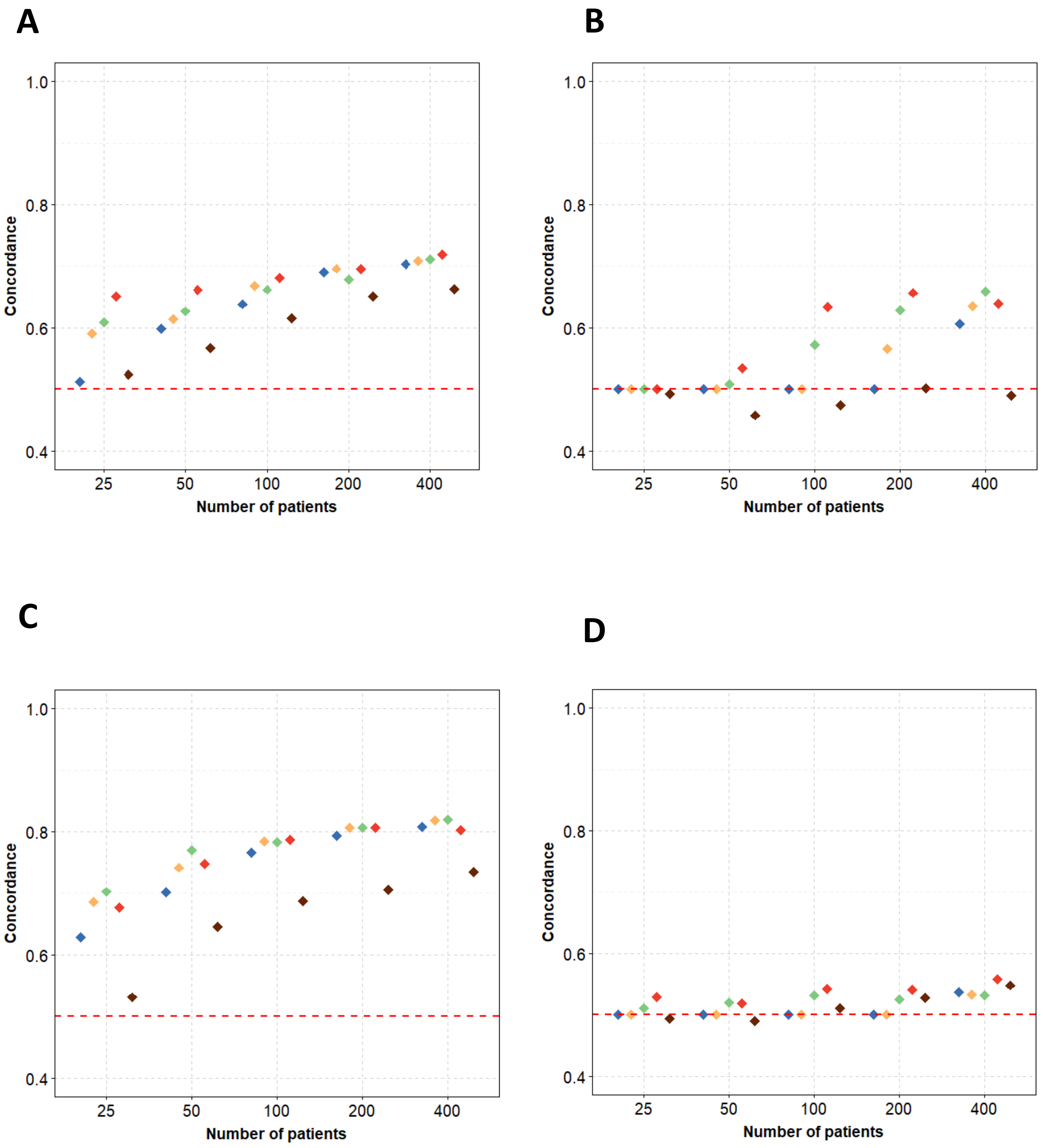


**Supplementary Fig. 10. Impact of the number of patients on the C-index computed with the genes with IQR ≥ 3 for the real TCGA dataset for kidney renal clear cell carcinoma (A), breast invasive carcinoma (B), brain lower grade glioma (C), and lung squamous cell carcinoma (D).** Medians of the C-indices as a function of the number of patients in the training dataset, for the lasso (blue), the elastic net (orange), the adaptive elastic net (green), the ridge (red), and the univariate Cox (dark red).
